# Prefrontal representations of retrospective spatial working memory in a rodent radial maze task

**DOI:** 10.1101/2024.10.10.617655

**Authors:** Joshua Paul Taliaferro, Lorenzo Posani, Julia Greenwald, Sean Lim, Josephine Cecelia McGowan, Elizabeth Pekarskaya, Clay Lacefield, Stefano Fusi, Christoph Kellendonk

## Abstract

Working memory is the cognitive capacity for temporarily holding information in mind for processing or use. It has been theorized to depend upon executive and mnemonic subcomponents, although the contextual mapping of these subcomponents is not complete. Perturbations of prefrontal cortex (PFC) delay activity disrupt spatial working memory performance in rodent tasks. However, recordings of unperturbed PFC delay activity do not consistently contain mnemonic representations of spatial information in these tasks, calling into question the role that mnemonic PFC representations play in freely-moving spatial working memory. We hypothesized that increasing task complexity might increase the likelihood of mnemonic PFC representation emergence. We therefore used an automated eight-arm radial maze to implement a novel match-to-sample rodent spatial working memory task with seven options on each trial, and recorded calcium activity in PFC neurons during task performance. We found that the delay-phase activity of PFC neurons indeed contained mnemonic representations of spatial information at the population level. These representations were retrospective rather than prospective, and—surprisingly—were more evident on error trials. Together with previous results, these observations suggest that in freely-moving spatial working memory tasks, PFC mnemonic representations emerge to empower deviation from a routine behavioral strategy.

**Significance Statement:** Prefrontal cortex (PFC) activity is necessary for optimal performance of freely-moving spatial working memory tasks in rodents. Despite this, PFC representations of retrospective actions or stimuli—one quintessential working memory hallmark—are only variably observed during task delays, complicating our understanding of the PFC’s role in spatial working memory. Here, we examine cellular-resolution PFC activity in a high-optionality match-to-sample radial maze task and find retrospective delay representations. Strikingly, these delay representations are more evident in error trials. This suggests that in the freely-moving context, explicit PFC representations of retrospective information support deviations from an entrained behavioral strategy, rather than equally supporting all spatial working memory-based behavior.

## Introduction

Working memory (WM), the mental capacity that enables temporary storage and manipulation of information, is foundational to myriad cognitive processes and daily activities. Consequently, WM deficits, such as those common with neuropsychiatric illness, can have catastrophic effects on an individual’s independent living and participation in larger society (Davidson and Stayner, 1997; Millan et al., 2012; Culbreth et al., 2021). While WM has been extensively studied, our knowledge about its neural correlates is still emerging. To facilitate the eventual development of treatments for WM deficits, a more comprehensive characterization of the neural basis of WM is necessary.

Although it has a long history (Locke, 1706; James, 2007), WM began receiving extensive formalized consideration around the middle of the 20^th^ century (Richardson et al., 1996; Cowan, 2014). Some of the resulting WM models, like that of Atkinson and Shiffrin <u>(1968)</u>, focused on information storage, defining WM as the system that held information on the order of seconds to minutes, between the millisecond-order storage of a sensory register and the days-to-years-order storage of long-term memory. Baddeley and Hitch <u>(1974)</u> later proposed that the WM system had multiple component types: temporary storage units and an executive function unit. The inclusion of an executive function component emphasized that WM enabled active manipulation of information, including by dictating which information entered storage, and by performing cognitive operations on stored information (Van Ede and Nobre, 2023).

The prefrontal cortex (PFC) has been considered a major hub of WM instantiation since nonhuman primates (NHPs) with PFC lesions showed deficits in two-choice delayed response WM tasks (Jacobsen, 1935; Jacobsen et al., 1936). Curiously, however, once neural activity could be recorded, some PFC neurons exhibited sustained activity during the task delays, but very few neurons (0-6%) exhibited delay activity that differentiated between the task cues ostensibly being held in WM (Fuster and Alexander, 1971; Niki, 1974). This suggested that this PFC delay activity was not primarily for storage of mnemonic representations. In contrast, when Funahashi, Bruce, and Goldman-Rakic <u>(1989)</u> created an eight-option, gaze-based WM task, more than 40% of recorded PFC neurons displayed delay activity with distinguishable mnemonic representations of cue information. The explanation might lie in the fact that the delays of Funahashi et al. 1989 are significantly shorter than those of some of the other articles. However, there could be a deeper reason: the comparative neural contributions of PFC activity to the executive and storage components of WM might depend heavily on behavioral context. Subsequent research has explored the intricacies of such neurobehavioral mappings (Meyers, 2018; Miller et al., 2018; Sreenivasan and D’Esposito, 2019; Buschman and Miller, 2022; Van Ede and Nobre, 2023).

In rodents, WM has often been studied in freely-moving spatial tasks with two choices, such as a T-maze delayed nonmatch to sample task. Notably, mnemonic PFC delay representations of retrospective actions are often absent in these tasks (Jung et al., 1998; Ito et al., 2015; Spellman et al., 2015; Bolkan et al., 2017; Böhm and Lee, 2020; Vogel et al., 2022), even though they are known to depend on PFC activity (Vogel et al., 2022). What role might mnemonic PFC representations play in these kinds of spatial WM tasks, if any? Given the NHP literature, a higher-optionality rodent task could help elucidate the context dependencies of the executive and representational components of the PFC’s role in freely-moving WM.

Accordingly, we created a new delayed match to sample spatial WM paradigm in an automated radial arm maze (RAM), inspired by both rodent RAM tasks (Olton and Samuelson, 1976; Saxe et al., 2007) and NHP tasks (Funahashi et al., 1989). We carried out cellular-resolution calcium imaging in the PFC of mice performing this novel task and observed mnemonic delay representations of retrospective spatial locations. Unexpectedly, however, these mnemonic representations were more evident on error trials. These findings suggest, therefore, that in freely-moving spatial WM tasks, mnemonic PFC representations preferentially emerge to help the animal change its behavioral strategy.

## Materials and Methods

### Animals

All animal studies were approved by the New York State Psychiatric Institute (NYSPI) Institutional Animal Care and Use Committee (IACUC) and the Columbia University IACUC. All experiments were carried out on C57BL/6J female mice purchased from Jackson Laboratory. A total of 10 mice were used. Mice were aged 8 weeks at the start of experiments and were housed under a reversed 12-h dark/12-h light cycle in a temperature-controlled environment (22 °C, humidity of 30–70%). All cages had running wheels. For initial training, mice were group housed with littermates (5 mice/cage). Mice with implanted GRIN lenses were individually housed. During behavioral training and testing, mice were food-restricted to no lower than 85% of their initial weight. Water was available ad libitum. After food restriction initiation, animals were habituated to the 20mg sweetened pellets (“sugar pellets”) used for behavioral rewards.

### Behavioral Procedures

All behavioral experiments were performed in the dark cycle.

### Radial Arm Maze

Behavioral experiments were conducted in a custom-built, automated 8-armed radial arm maze. The maze surfaces were made of gray acrylic plates (Canal Plastics), elevated on a steel frame (80/20 Incorporated). It featured an open, octagonal center, 80 cm in diameter, with a 30 cm arm coming off of each side. Each of the arms could be flexibly closed off from the center by a pneumatic door that rose vertically from the maze floor, under control of a solenoid array (Noldus). At the end of each arm was an automated 20-mg pellet dispenser (Med Associates). 24 LED strips (LEDGlow) symmetrically illuminated the maze, with 16 along the interarm walls of the maze center, and 8 at the ends of the arms.

The maze surface was 85 cm above the ground, and the entire maze was enclosed in floor-to-ceiling microscopy-grade blackout curtains on a custom-designed PVC pipe (McMaster-Carr) apparatus to help control auditory, visual, and olfactory cue exposure. A wide-angle HD webcam (Logitech) was secured to the PVC frame, 115 cm above the maze surface. Paper cutouts of geometric visual cues were asymmetrically arranged on the inside of this curtain enclosure. A white noise machine (LectroSound) and a 0.18 – 20 kHz frequency response speaker (JBL) sat below the maze. The maze doors, pellet dispensers, and cue LEDs were controlled locally by an Arduino (Arduino Inc.), which interfaced with a first-generation Any-maze Interface I/O box (Stoelting Co.). This latter device was connected to a computer (Hewlett Packard) with Any-maze software (Stoelting Co.), where a custom script enabled control of maze operations, dynamically dependent on the animal’s position in the maze, as captured by a convergently-connected webcam. For calcium imaging experiments, this computer was also connected via the Any-maze interface to the miniature microscope data acquisition box (Inscopix), through which calcium imaging data and behavioral data could be synchronized on a frame-to-frame level.

### Delayed Match to Sample Working Memory Task

The training and task were conducted in the radial arm maze. One mouse was run at a time. 8 mice were trained in the following manner:

### Habituation

Mice were given 3 days to habituate to the maze with all arms open. They could freely explore the maze for 20 minutes. Two reward pellets (dustless precision pellets, Bio-Serv) were available at the end of each arm and in the very center of the maze.

### Shaping

For 3 consecutive days, mice were placed in the start arm (arm 1), with all doors closed. Each trial then consisted of the opening of the door to arm 1 and to one other randomly selected arm, allowing the mouse to visit the non-start arm. When the 80% of the mouse’s body entered the non-start arm, as automatically determined by a computer vision contingency, a single reward pellet was dispensed at the end of the arm. Upon the mouse’s return to the start arm, another reward pellet was dispensed, both doors would close, and the trial was completed. For the next trial, a different randomly selected arm would become the one open arm, and the procedure would repeat. Mice had 30 minutes to complete as many trials as possible.

### Further training

Mice were then iteratively trained in versions of the task that increasingly approximated the full task. These training versions progressed in terms of both increasing number of incorrect arms open during the choice phase, increasing their proximity to the correct arm, and decreasing the number of rewards available per trial. Each version followed the basic structure of trials with four phases: Intertrial Interval (ITI), Sample, Delay, and Choice. In all training stages, the ITI duration was 28s, and the delay duration was 2s. The mice had 45 minutes to complete as many trials as possible.

The training progression occurred as follows, with the same basic structure, but varying arm openings:

| Training Version | Sample Arm Possibility | Choice Arm Possibilities |
| --- | --- | --- |
| Avoid consistent distractors | One from 3-7 | Sample arm + 2 and 8 |
| Avoid variable distractors | One from 4-6 | All arms 4-6 |
| Avoid variable distractors II | One from 3-7 | All arms 3-7 |
| Full task | One from 3-7 | All arms 2-8 |

Consistent distractors: during the choice phase, arms 2 and 8 are open on every trial, but are never rewarded, and are therefore distractors.

Variable distractors: during the choice phase, all of the arms under “Choice Arm Possibilities” are open on each trial. Whichever arms were not the sample arm are not rewarded, and are therefore distractors.

The mouse began each behavioral session confined to Arm 1, deemed the start arm. Each trial then proceeded through the following phases:

### Sample

An arm from the possible sample arms was randomly selected to be that trial’s sample arm. When the trial started, the doors to the start arm and the sample arm were both lowered, and the mouse could visit the sample arm. As soon as the mouse entered the sample arm, a 19 KHz tone was played for 0.1 seconds. No reward was dispensed. The animal had to then return to the start arm, at which point both doors closed, ending the Sample phase.

### Delay

During the Delay phase, the animal was confined to the start arm for the delay duration (2s). No reward was dispensed.

### Choice

When the Choice phase began, the doors to all possible choice arms opened, and the animal was free to visit any open arm. If the animal returned to the sample arm (“correct” choice), as soon as they entered it, a 19 KHz tone was played for 1s, and a sugar pellet (Dustless Precision Pellets, 20 mg, Bio-Serv) was dispensed at the end of the arm. If, instead, the animal visited any other arm (“incorrect” choice), as soon as they entered their chosen arm, white noise was played for 1s, and no reward was dispensed. In either case, as soon as the animal made a choice, the doors to all other arms (besides the start arm) closed. The animal then had to return to the start arm, at which point both of the still-open doors closed. If the animal made the “correct” choice, a second sugar pellet was dispensed upon their return to the start arm; if “incorrect”, no reward was dispensed.

### Intertrial Interval (ITI)

During the ITI phase, the animal was confined to the start arm for the ITI duration (28s). After this a new trial began. The session ended when 45 minutes had elapsed.

### Shaping Reminder Trials

If the mouse got the trial correct, the sample arm for the subsequent trial would change to another pseudorandomly selected arm from the sample arm possibilities for that training stage. If, instead, the mouse got the trial incorrect, the sample arm would stay the same for the subsequent trial. If the mouse got three consecutive trials incorrect with the same sample arm, the subsequent trial would be a shaping trial, in which the sample phase and choice phase both only had the same single non-start arm open, thus obligating the mouse to get the trial “correct.” The sample arm of the subsequent trial would then become a pseudorandomly selected arm from the sample arm possibilities for that training stage. After training, mice were given ad libitum food and water until their weight recovered to baseline.

### Surgical Procedures

Mice were first anesthetized with isoflurane and head-fixed in a stereotactic apparatus (Kopf). An AAV9 expressing GCaMP6f under a CaM Kinase II promoter (pENN.AAV9.CamKII.GCamp6f.WPRE.SV40, titer ≥ 1×10¹³ vg/mL, Addgene) was then injected bilaterally into the medial prefrontal cortex (mPFC), at a volume of 600 nL per side, at three sites on the dorsal-ventral axis. Injection coordinates were +1.98 AP, −2.3 /-2.15/-2.0 DV skull, ±0.35 ML. Mice were also injected bilaterally with virus (AAV5-CaMKIIa-eNpHR3.0-mCherry, Addgene) in the mediodorsal thalamus, but no optogenetics were performed for the experiments in this paper.

Additionally, mice were also unilaterally implanted with a gradient index (GRIN) lens (6.1 mm length, 0.5 mm diameter, Inscopix #1050-002211) in the right mPFC, at coordinates +1.98 AP, −2.0 DV skull, +0.35 ML, which was secured to the skull with dental cement (Stoelting Dental Cement, Fischer Sci. #10-000-786). Two weeks later, mice were fit with a baseplate (Inscopix #1050-004638) to allow for attachment of the miniature microscope. This was secured to the GRIN lens and skull with dental cement.

One mouse died immediately after surgery. Two other mice, without the behavioral training history described above, underwent surgery in the described manner and were subsequently trained alongside the other seven mice in post-surgical re-training. All nine mice were then imaged.

### Calcium Imaging Experiments

Once recovered from surgery, mice were gradually food-restricted to no less than 85% of their baseline body weight, and were retrained on the delayed match to sample task until the group mean performance reached 50% correct. Mice were then imaged every other day, with five mice imaged on one day, and four mice on the other.

For the imaging days, a mouse was anesthetized with isoflurane and a miniature microscope was attached to their baseplate. After 30 minutes to allow for recovery from anesthesia, the mouse was run on the full task, with synchronized behavioral video and calcium imaging data coordinated by the computational setup described above, with collection parameters described below. Calcium activity was recorded for the duration of the experimental session (forty-five minutes). The mouse was then briefly anesthetized with isoflurane for miniscope removal.

### Data Acquisition and Preprocessing

Neural recordings were collected at 20 Hz exposure (gain 1.0–2.0, exposure time 50 ms, LED power 0.2-1 mW) using Inscopix data acquisition software. Videos were then spatially downsampled by a factor of 4, cropped to the lens, and exported to tiff stacks using Inscopix Data Processing (IDP) software. Single cell fluorescence activity traces were extracted from background using a Python implementation of Caiman (Giovannucci et al., 2019), running NoRMCorre (Pnevmatikakis and Giovannucci, 2017) for motion correction, and constrained non-negative matrix factorization for endoscopic data (CNMFe) for signal extraction <u>(Zhou et al., 2018)</u>. Data matrices were imported as .hdf5 files to MATLAB for further processing.

### Histology

At the end of experimentation, mice were transcardially perfused with PBS followed by 4% PFA. To preserve lens tracts, brain tissue remained in 4% PFA for 48 hours, before being transferred to PBS. Fixed tissue was then sectioned (50 μM) orthogonally to the AP axis using a vibratome (Leica), and mounted on slides with Vectashield mounting medium containing DAPI (Vector Labs). Direct fluorescence of GCaMP was then examined under an epifluorescent microscope (Zeiss) to assess the extent of viral spread and expression pattern. Locations of recording site were confirmed with visualization under DAPI. No mice were excluded from analyses due to failed lens targeting.

### Experimental Design and Statistical Analyses

#### Data Utilized

Two calcium imaging sessions of 45 minutes each were performed for nine mice. A total of 3979 neurons were imaged across these 18 mouse-sessions. For single cell analyses, neurons are considered individually. For population analyses, neurons are considered collectively as a single pseudosimultaneously recorded population. To mitigate the effects of photobleaching, only trials that were completed in the first 25 minutes of each session were used for analysis, unless otherwise noted.

#### Single-Cell Selectivity Analyses

Events inferred from calcium transient activity were used for single-cell analyses. The event rate was estimated from calcium transients (Pnevmatikakis et al., 2016). Though this method cannot definitively identify true firing rates, it does provide a proxy for each neuron’s relative firing rate.

Assessment of spatial information content was performed similarly to that in Stefanini et al. <u>(2020)</u>. Briefly, given a cell *i*, the mutual information score was calculated by computing the entropy of the mean activity rate across trial type (either sampled arm, or chosen arm):

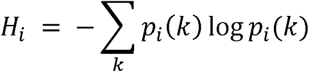

Where *p_i_(k)* represents the mean activity rate of cell *i* given the arm label *k*, normalized such that *Σ_k_p_i_(k) × 1*. For each cell, a null distribution was then generated by shuffling the arm label across trials, making sure that each trial was coherently labeled with one arm type only. The mutual information score for the data was then z-scored against the null distribution to obtain the values shown in the figures. These analyses were performed during the two seconds in the middle of the delay phase.

#### Concurrent Population Activity Analyses

All of these analyses were performed on data from the delay phase. Both “correct” and “incorrect” trials were included. Only mouse-sessions in which at least two instances of each trial type were completed by the time cutoff were included, because this would allow at least one instance for training the decoders, and one for testing the decoders. This resulted in 13/18 mouse-sessions being included to build the single pseudosimultaneously recorded population (pseudopopulation) (Meyers et al., 2008). This approach assumes similar neural activity statistics across all mouse-sessions, and partially disrupts the contributions of noise correlations (that would otherwise be present if neurons were in fact all simultaneously recorded) to the population code (Abbott and Dayan, 1999). For these analyses, CNMFe-extracted fluorescence traces of cells were used, after undergoing soft-normalization to prevent the signal from being dominated by a few highly active cells, while still allowing for the possibility of higher activity conveying some useful information (Churchland et al., 2012; Seely et al., 2016; Russo et al., 2018, 2020). The equation for soft-normalization was *response:= response / (range(response) + 5)*.

#### Decoding

To search for delay representations of spatial information, we used a support vector machine (SVM) classifier with a linear kernel to perform decoding on the pseudopopulation data. Neural activity was averaged within a trial to reduce the pseudopopulation activity matrix from three-dimensional *trials x neurons x timepoints* to the two-dimensional *trials x neurons*. Trials were labeled based on either the arm the animal had visited during the sample phase (for decoding retrospective spatial information), or the arm the animal would later choose to visit during the choice phase (for decoding prospective spatial information). These labels were identical in all “correct” trials, but diverged in “incorrect” trials. Trials were split randomly into training and testing sets in a 75:25 manner; the SVM classifier was trained on trials from the training set, and then tested on trials from the testing set. This random subsampling cross-validation process was repeated 25 times; decoder accuracy was reported as the average fraction of testing trials that were correctly classified. To evaluate the statistical significance of decoding spatial information, we built null models by shuffling the trial labels in the training set and repeating the whole procedure 100 times to obtain a null model distribution of decoding accuracy for spatial information (retrospective or prospective). The average decoding performance on the unshuffled data was z-scored against this null distribution and given a corresponding p-value. All decoding analyses were implemented with custom scripts built on the Statistics and Machine Learning Toolbox in Matlab (The MathWorks, Inc.).

#### Importance Index and Gini Coefficient

Given that multinomial SVM classifiers are amalgamations of multiple binary SVM classifiers, assessing the contributions of each cell to decoding requires some manner of combining decoder weights across classifiers. We used the importance index from (Stefanini et al., 2020), which is, for each cell, a normalized sum of decoder weights. From the value of this for each cell, we could build a distribution of importance indices across the population. To then assess how distributed the population code is across cells, we used the Gini coefficient (Gini, 1912; Ceriani and Verme, 2012), which is a measure of statistical dispersion used to characterize distributional inequality.

#### State Space Analyses

The average representation of each trial type in state space was plotted for both individual exemplar cells and for the first three principal components (Figure 4). For the individual cells, three random cells were selected, and the activity of each cell over the delay was averaged across each trial type, before being plotted in the neural activity space. For the first three principal components, principal component analysis was performed on the full matrix of neural activity of the pseudopopulation during all trials to determine the dimensions explaining most of the population variance. From there, the activity along each of the first three principal components was averaged by trial type and then plotted in the principal component space. The first three principal components explained 24.7% of the variance.

#### Cross-temporal Arm Decoding

To assess the temporal stability of representations, cross-temporal decoding was performed during the delay for a five-dimensional approximation of the data from the first five principal components. A linear SVM classifier was trained on neural activity from one delay timepoint (one slice of the third dimension of the *trials x neurons x timepoints* data matrix), labeled by that trial’s sample arm, and was tested individually on the neural activity at every delay timepoint (including the training timepoint). Decoder accuracy was calculated as the fraction of trials with a correctly classified sample arm at each testing timepoint. The data matrix was a trials x principal components x timepoints matrix created by performing PCA on the original matrix, and only the first five principal components were used (which explained 33.4% of the variance).

#### Cross-arm Temporal Decoding

To further assess the temporal consistency of representations across trial types, cross-arm temporal decoding was performed during the delay for the full data set. Each timepoint of the *trials x neurons x timepoints* data matrix was labeled as either early or late, depending on whether it came in the first half or second half of the delay. Using data from a single trial type, a linear SVM classifier was trained to temporally classify neural activity as either early or late within the delay. This classifier was then tested on neural activity from each of the different trial types, one at a time; the accuracy of the classifier with different combinations of training data and testing data is shown in Figure 4.

## Results

### A novel task in an automated radial arm maze offers more spatial optionality and complexity

Inspired by the increase from two to eight task options that had proven so illuminating in the nonhuman primate (NHP) domain (Funahashi et al., 1989), we adapted existent spatial working memory (WM) tasks for mice to similarly increase task optionality while decreasing the possibility of subversion of WM requirements (Chudasama and Muir, 1997). To begin, we built an automated radial arm maze (RAM), with eight arms attached to a large octagonal center (Figure 1A, B). An Arduino-based system allowed the maze’s pneumatic doors and pellet dispensers to be dynamically controlled by the animal’s position (see *Materials and Methods*). We then drew upon a rich history of radial arm maze tasks (Olton and Samuelson, 1976; Olton et al., 1977; Olton, 1987; Packard et al., 1989; Brown et al., 1993; Jarrard, 1993; Floresco et al., 1999; Redish, 1999; Dudchenko, 2004; Wenk, 2004; Saxe et al., 2007; Foreman and Ermakova, 2013; Missaire et al., 2017) to design a new task (Figure 1C). On each trial of this Delayed Match to Sample (DMS) task, mice would visit one arm, randomly selected from arms 3-7, during the Sample phase, return to the start arm, and then, after the Delay, would choose from seven arms (2-8) to visit, only getting the trial correct if they returned to that trial’s sample arm.

**Figure 1.**
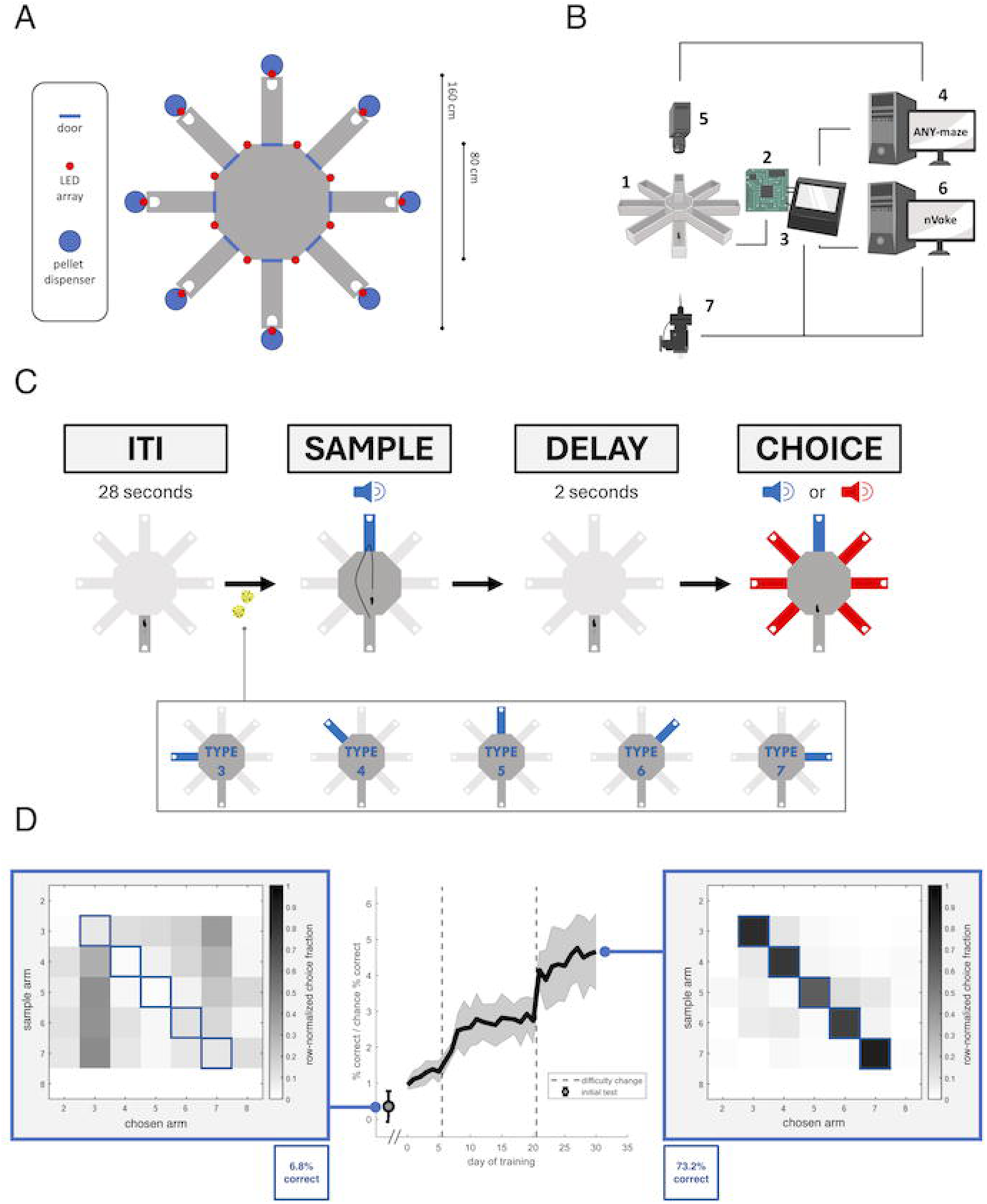
Mice can acquire and stably execute a novel multi-alternative rodent spatial working memory task. A. Design of the maze surface. Doors are pneumatic doors that rise from below to prevent movement into/from each arm. LEDS are linear light arrays, symmetrically arranged for even lighting. Automated pellet dispensers (PD) provide 20-mg sugar pellets as reward. B. Maze connectivity. The doors and pellet dispensers of the maze (1) are controlled locally by an Arduino (2), which connects through an interface box (3) to two computers: one (5) with Any-maze behavioral tracking software, which takes input from the overhead camera (6) to allow for maze control based on the animal’s position; and one (4) running Inscopix software for performing cellular-resolution calcium imaging via the miniscope (7). C. The delayed match to sample task. On each trial, the animal begins in the start arm. When the trial starts, a sample arm is pseudorandomly chosen from arms 3-7, and the animal must visit that arm before returning to the start arm. After a delay of 2s, all arms open, and the animal can visit any arm, but will only get a reward if they return to the sample arm. In between each trial, there is a fixed intertrial interval of 28s. If the animal gets the trial correct, the subsequent trial has a new pseudorandomly-selected sample arm from arms 3-7 (with no consecutive sample arm repeats). If the animal gets the trial incorrect, the subsequent trial repeats the same sample arm. Speakers represent different auditory cues (see *Materials and* Methods). D. Acquisition of the task. The middle panel shows the group-averaged ratio of actual percentage correct over chance percentage correct (14.3%) as the animals were trained in progressively harder versions of the task. Dashed vertical lines indicate where task difficulty was increased until the full task was used. Error bars are SEM for n = 8 mice. The leftmost datapoint of this graph is the first day of exposure to the task, immediately after shaping. This corresponds to the left heatmap, which shows the choices the animals made across all trials and mice on that first day. Each row represents that trial’s sample arm, while each column represents the potential arms animals could choose. Entries along the diagonal are therefore correct choices (matches to sample). On this first day, animals tended to visit arms 3 and 7, no matter the trial type, and accordingly got only 6.8% of trials correct (0.48x chance performance). In contrast, the heatmap on the right shows collective group performance on the last day of training (corresponding to the rightmost data of the middle panel). By then, animals had acquired the task, getting 73.2% of trials correct (5.1x chance), and showing diffuse and non-perseverative errors. In both heatmaps, values are number of trials, normalized by the total number of trials in each row.

Unlike in classic versions of the RAM spatial working memory task, mice can complete several dozen trials per session in this task. Unlike other higher-throughput RAM spatial working memory tasks (Saxe et al., 2007; Missaire et al., 2017), each trial here has seven options, dramatically lowering the chance floor. And unlike another higher-throughput, higher-optionality win-stay RAM spatial working memory task where the animal spends the delay in the maze center (Jarrard, 1993), the use of a start arm prevents the mouse from subverting the working memory requirement by simply sitting in front of that trial’s sample arm during the delay. Additionally, automation decreases the likelihood of the experimenter affecting data collection (Bohlen et al., 2014; Nigri et al., 2022). All told, the task differs from its predecessors in ways that facilitate the further clarification of neurophysiological working memory correlates.

Over the course of training, mice acquired the task, progressing from performing below chance to performing more than five times chance levels (Figure 1D, center). While on day 0, mice got 6.8% of trials correct, with errors mainly to the arms near the start arm (Figure 1D, left), by the last day of training assessment on the task, mice got 73.2% of trials correct, with errors that tended to be close to the correct arm, and were mostly diffuse instead of perseverative (Figure 1D, right).

### Cellular-resolution calcium imaging of the PFC in the automated radial arm maze

To investigate the contribution of the PFC to spatial working memory, we measured PFC activity at cellular resolution with miniature microscopes (Flusberg et al., 2008) during task performance. Trained mice were injected with an Adeno Associated Virus (AAV) expressing GCaMP6f under a CamKIIalpha promoter to target excitatory projection neurons, and implanted in the right PFC with a 500-um gradient index lens (Figure 2A-C, *Materials and Methods*). After post-surgical recovery and retraining, PFC pyramidal cells were imaged as mice performed the behavioral task. Figure 2C shows the field of view of PFC cells with GcaMP6f-mediated fluorescence in a representative mouse, while Figure 2D shows exemplar task-evoked calcium activity traces for individual cells. All together, we imaged n = 3979 cells from 9 mice over 18 sessions (mostly in the prelimbic subregion of the PFC), which for subsequent analyses are considered as a single pseudosimultaneously-recorded population (Spellman et al., 2015; Stefanini et al., 2020)

**Figure 2.**
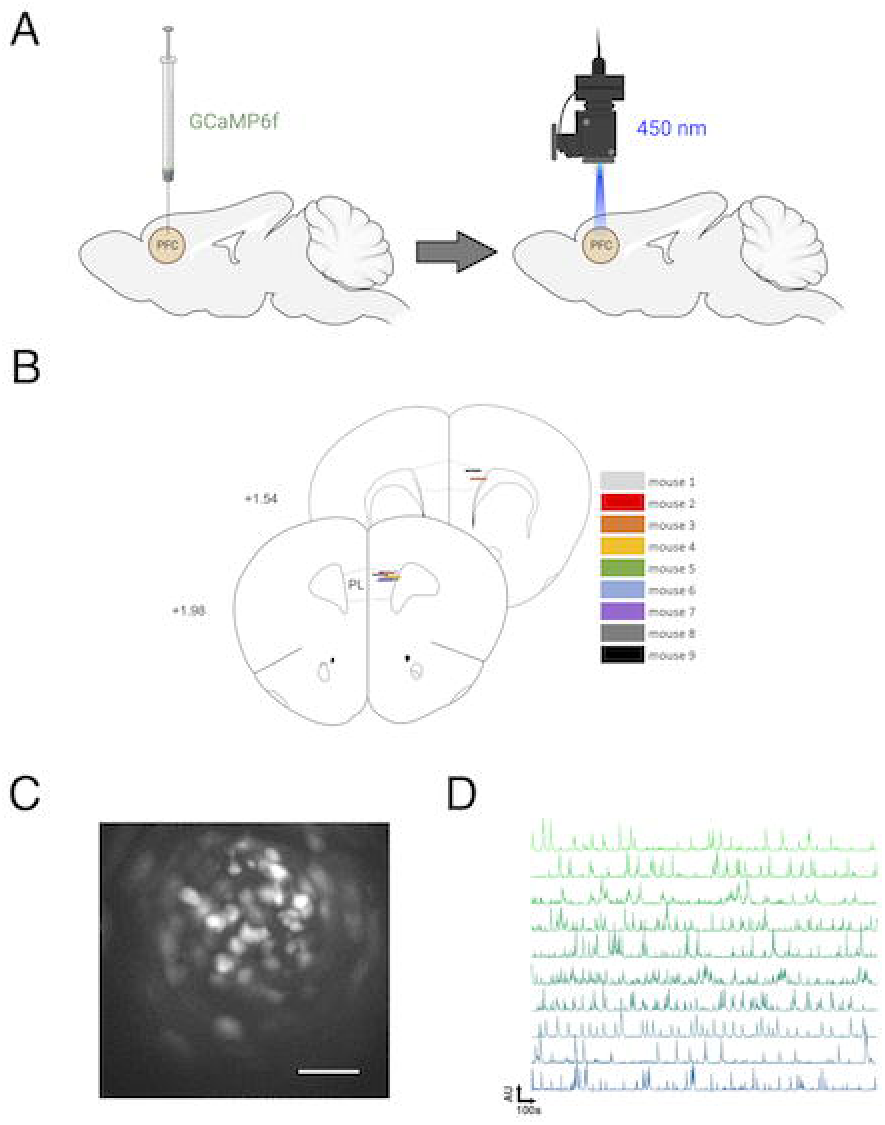
Microendoscopy allows for cellular-resolution observation of population activity in freely-moving mice during task. A. Simplified surgical scheme. Mice were injected with a virus expressing GCaMP6f under a CamKII promoter in the right medial prefrontal cortex (medical PFC). After implant of a GRIN lens and baseplate attachment (not shown), calcium imaging could be performed using a miniaturized microscope. B. Location of GRIN lenses for n = 9 mice. C. Representative field of view from one imaging session. D. Representative extracted single-cell calcium transients.

### Minimal delay representations of spatial information are evident at the single-cell level

Given the robust history of single-cell representations of WM content (Goldman-Rakic, 1995), and the evident heterogeneity of single-cell delay responses to spatial stimuli in our recordings (Figure 3A,B), we first sought to systematically characterize single-cell spatial representations in this task. To do so, we calculated the normalized mutual information between spatial location (the sample arm), and the cell activity during the two second delay period, before comparing with a null distribution generated from the same dataset with shuffled arm labels (similar to Stefanini et al., 2020).

**Figure 3.**
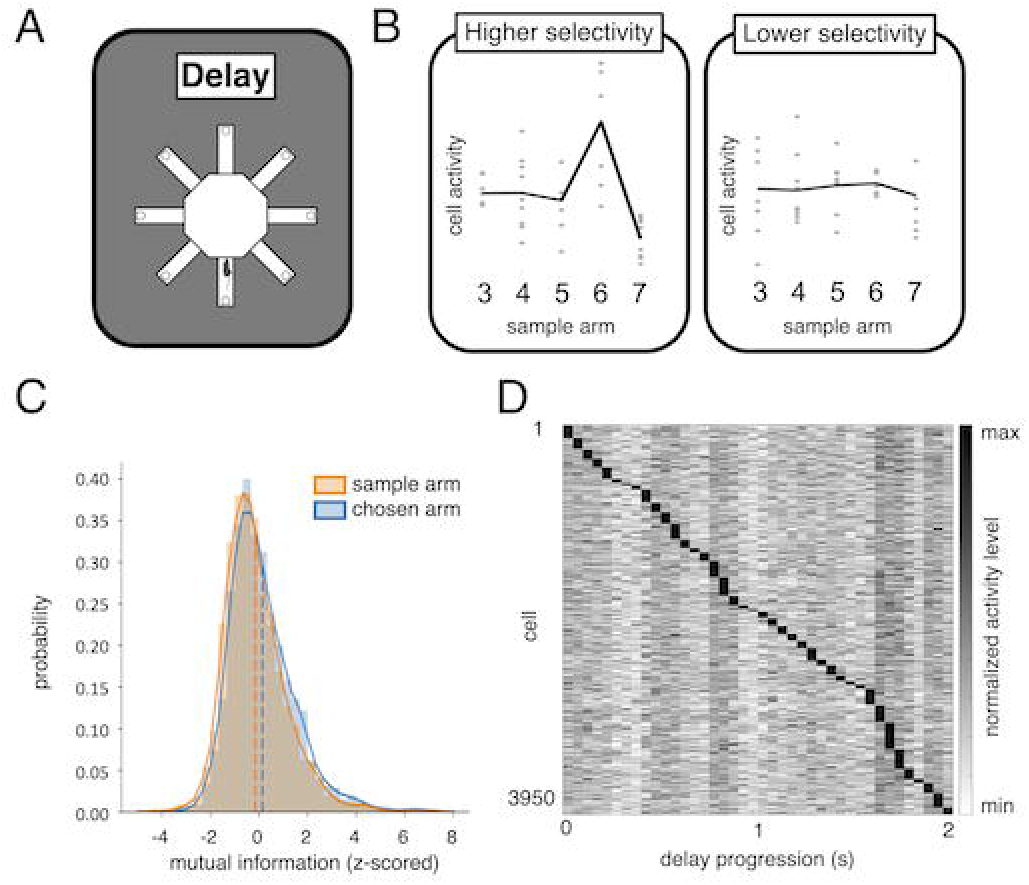
Single cell delay activity contains few cells with significant representations of spatial information. A. GCaMP activity of individual neurons was assessed during the two second delay period, after the animal had returned from visiting the sample arm. B. Cells differed in the extent to which their average delay activity was affected by the sample arm they had just visited. C. Single-cell spatial information. Spatial information was calculated as the entropy of normalized mean firing rates across the different arm labels, z-scored against a null distribution obtained by shuffling the arm labels (see *Materials and Methods*). Using a significance cutoff of p = 0.001 for deviation from the distribution mean, 3.1% and 1.8% of cells were selective for retrospective and prospective spatial information, respectively, D. Temporal pattern of peak activity. Average delay activity traces were calculated for each cell, and normalized across cells. These traces were then plotted in order of the earliness of the respective peak of that cell’s average, normalized delay activity trace. Cell activity peaks span the delay. These analyses were all performed on n = 3979 cells.

Studies featuring freely-moving, two-choice T-maze tasks tend to find no single cells selective for retrospective or prospective spatial information during the delay phase (Bolkan et al., 2017; Vogel et al., 2022). In a similar manner, we found only a small proportion of retrospective or prospective spatially-selective cells during the delay phase (Figure 3C, retrospective: 3.1% of cells, prospective: 1.8% of cells). In further accordance with these studies, we also found that single cells tended to be transiently active during short windows (50-150ms, Figure 3D), rather than exhibiting the sustained, delay-spanning activity changes commonly observed in head-fixed NHP WM tasks (Goldman-Rakic, 1995; Wang, 2001). Altogether, then, the increased optionality of this spatial working memory task seems to have had little effect on single-cell activity patterns or representations of spatial information during the delay.

### Delay representations of spatial information are evident in heterogeneous population activity

To further characterize PFC spatial representations, we next considered how the activity of individual cells might be coordinated to collectively contain spatial information not evident when considering the activity of each cell individually. Here, we focused on population activity during the delay period of the task, in which the mouse is presumably using WM to maintain a representation of the arm visited during that trial’s sample phase.

To assess the extent to which population representations of spatial information are distinguishable for individual trials, we performed a decoding analysis. We used a simple support vector machine (SVM) decoder with a linear kernel (*Materials and Methods*), based on the notion that downstream neurons could feasibly implement such a decoder by changing input weights. We found that the decoder performed beyond chance at identifying the trial’s sample arm (Figure 4B, average 49.7% accuracy on held-out testing trials across random subsampling cross-validations), significantly beyond the performance on data with shuffled trial labels (average 19.4% accuracy, p value of difference p = 0.001 via z-scoring). Notably, this performance was lost when decoding animal’s forthcoming choice (Figure 4A, right, average 18.7% accuracy), which was statistically indistinguishable from the shuffled data set decoding (average 20.6% accuracy, p value of difference p = 0.42 via z-scoring). Together, these data suggest that the delay population activity of the PFC in our task does contain retrospective—but not prospective—spatial information.

**Figure 4.**
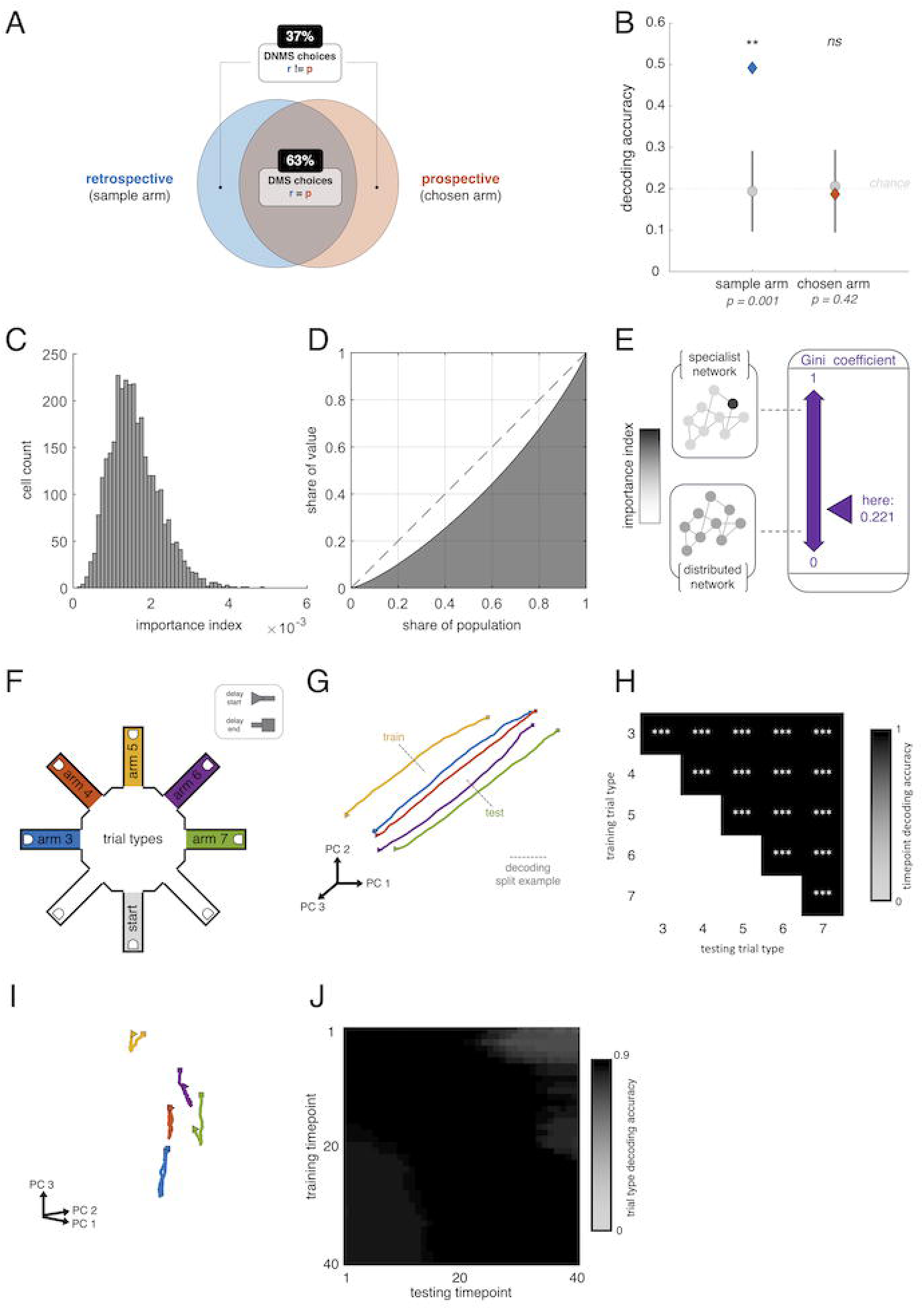
Distributed representations of retrospective spatial information are evident in prefrontal cortex population delay. A. Under this delayed match-to-sample (DMS) task design, correct trials have identical retrospective and prospective spatial information labels. Incorrect trials, on which animal employs a delayed nonmatch-to-sample (DNMS) behavior, have divergent labels. B. Retrospective and prospective decoding analysis during the delay. A support vector machine (SVM) with a linear kernel was trained to classify trials using delay activity of a pseudopopulation of the cells of all mouse-sessions with at least two instances of each condition (3180 cells from 13 mouse-sessions). Decoding of retrospective spatial information was possible with an accuracy exceeding that of a null model (p = 0.001), while decoding of prospective spatial information was indistinguishable from the null model (p = 0.42). Error bars are standard deviation. C. Distributed decoder contributions. The contribution of each cell to decoding was quantified using the importance index (Stefanini et al., 2020). The distribution of this value for all included cells is shown in the histogram. D. The Lorenz curve shows what proportion of cells shares what proportion of cumulative decoder importance, and is used to calculate the Gini coefficient for this distribution. E. The Gini coefficient ranges from 0 to 1, with lower values representing greater equality of the metric across the population, and higher values representing greater inequality of the metric across the population. Here, the Gini coefficient for the distribution of importance index across cells is 0.221, which suggests a rather distributed representation. F. Trial types are classified by the sample arm visited before the delay. Displayed trajectories begin at the arrow (▷), and end at the square (□). G. Exemplar neural activity space trajectories, view 1. The trajectory of average activity for each trial type was plotted in the state space, as defined by the first three principal components resulting from performing principal component analysis on the matrix of pseudopopulation activity data from the delay. Each axis is one principal component. Each trajectory represents the temporal evolution of average population activity over the delay for one trial type. H. Cross-arm temporal decoding. Population activity data were binarily labeled by time during the delay (first half vs. second half) as well as trial sample arm. A linear SVM was trained on temporally labeled data from trials with one sample arm, and tested for temporal labels on trials with another sample arm, for all ten possible combinations of sample arm. Decoder accuracy greatly exceeded chance for all combinations (decoding accuracy ≥ 0.99 and *p* < 0.001 for every combination), supporting the temporal alignment of state space trajectories observed in G. I. Exemplar neural activity space trajectories, view 2. As in G, each trajectory represents the temporal evolution of average population activity over the delay for one trial type. From this view, trajectories appear to somewhat recapitulate maze geometry, with a skew. J. Cross-temporal arm decoding for lower-dimensional subspace. Cross temporal decoding was performed by training a linear SVM on neural activity data from one delay timepoint, and testing it on neural activity data from all delay timepoints. Only the neural activity captured in the first five principal components was used. Decoder accuracy was defined by the proportion of testing trials that were correctly classified. The presence of many high off-diagonal accuracy values is consistent with a population code with significant time invariance.

This combination of decoding results was somewhat surprising given the task design. A match-to-sample task has identical retrospective and prospective spatial information on all trials that the animal gets correct, and divergent information on all trials that the animal gets incorrect (Figure 4A). Consequently, if correct trials were primarily driving the mnemonic PFC delay representations, one would expect retrospective and prospective decoding accuracy to be similar, if not equal. If, instead, error trials were primarily driving the mnemonic PFC delay representations, one would expect retrospective and prospective decoding accuracy to be quite distinct. Our data are consistent with the second scenario (Figure 4B). Curiously, this combination suggests that there was a stronger delay representation of the just-visited sample arm on trials in which the animal would subsequently avoid returning to that arm. Unexpectedly, then, mnemonic PFC delay representations seem to preferentially emerge to support a deviation from the dominant behavioral strategy under our task conditions.

From this initial foundation, we sought to further characterize the nature of these mnemonic PFC delay representations. Population-level representations can be driven by a few highly selective cells, or instead carried broadly across the population. To better understand how distributed these population-level representations were, we examined the weights assigned to each cell’s activity for classification by the decoder. Given that multiclass classification with SVMs requires stitching together multiple binary SVM classifiers, we used the decoder importance index (Stefanini et al., 2020) to characterize the comparative contribution of each cell to decoding across classifiers. Figure 4C shows a histogram of these importance indices. We then calculated a distributional Gini coefficient, which ranges from 0 for perfect equality, to 1 for perfect inequality (Gini, 1912; Stefanini et al., 2020). Here, the importance index Gini coefficient was 0.22 (Figure 4D, E), suggesting that the population of cells shows a tendency toward equality in importance to decoding sample arm. Thus, the retrospective spatial information representations during delays in this task seem to be carried by a heterogeneous, distributed population code.

### Delay representations of spatial information are evident in a stable low-dimensional activity subspace

How could a downstream region stably read out spatial information from such a heterogeneous population code with dynamic individual cell activity patterns? Abundant experimental work (Stopfer et al., 2003; Sadtler et al., 2014; Murray et al., 2017; Spaak et al., 2017; Parthasarathy et al., 2019; Cueva et al., 2020; Sapountzis et al., 2022; Voitov and Mrsic-Flogel, 2022) and modeling (Druckmann and Chklovskii, 2012; Shenoy et al., 2013; Gallego et al., 2017; Langdon et al., 2023) suggest that heterogeneous population codes often contain a low-dimensional subspace from which time-invariant information can be stably read out. To assess for the presence of such a low-dimensional representation in our data, we performed dimensionality reduction, via principal component analysis, on the matrix of delay activity of all cells across all trials (*Materials and Methods*). Visualizing the average activity of the different trial types in the principal component (PC) space could give us an intuition about the temporal stability we might expect to later assess more formally. Considering the first three principal components (which explain 24.7% of variance), we observed that although average trial type trajectories were not stationary (Figure 4G), their movement across the delay was largely parallel and non-overlapping, allowing them to appear stationary and steadily separable from a particular field of view in the principal component space (Figure 4I). And in fact, with similarities to other work (Murray et al., 2017), the arrangement of these trajectories in PC space recapitulated the maze geometry, with some compressive contortion (Figure 4F vs Figure 4I).

To assess the concordance of temporal trajectories across trial types, which seemed reasonably apparent in the state space visualizations of Figure 4G and 4I, we used a linear decoder to perform cross-sample arm decoding of time. Specifically, a linear SVM trained on binarized temporal labels from trials with one sample arm was tested on temporal labels from trials with the other sample arms. Temporal information could be read out significantly beyond chance across all pairwise combinations of sample arms used for training and testing (Figure 4H). In an analogous and complementary manner, to assess the temporal stability of population representations of spatial information during the delay, we performed cross-temporal decoding of the sample arm, in which a decoder is trained on population activity of a single timepoint, and then tested for its capacity to classify using population activity at every other timepoint (Stokes et al., 2013; Spaak et al., 2017; Meyers, 2018; Cueva et al., 2020). Training a decoder on the dimensionality-reduced population activity (first 5 PCs, explaining 33.4% of variance) at one delay timepoint led to solid performance in classifying the sample arm at many other delay timepoints (Figure 4J), consistent with the existence of a stable low-dimensional subspace. Together, the robust results of both the cross-temporal sample arm decoding and the cross-sample arm temporal decoding suggest that a downstream region could use stationary input weights to stably read out spatial information from a low-dimensional subspace of PFC activity throughout the delay.

## Discussion

### A high-optionality, freely-moving spatial working memory task evokes mnemonic delay representations in the prefrontal cortex at the population level (PFC)

Here we’ve introduced a novel rodent spatial working memory task (Figure 1). Building upon the rich history of radial arm maze (RAM) tasks, we developed a delayed match to sample task, in which the mouse must select the correct answer from seven options on each trial. We showed that mice can successfully acquire the task, performing well beyond chance (Figure 1).

We then recorded single-cell calcium activity in the prefrontal cortex (PFC) during the task with a miniaturized microendoscope (Figure 2) and found that, despite there being minimal PFC delay representations of retrospective spatial information at the single-cell level (Figure 3), there were such representations at the population level (Figure 4). We revealed that these representations are broadly distributed across the recorded cells, yet also contain a low-dimensional subspace with time-invariant retrospective stimulus information.

The combination of a heterogeneous, distributed activity profile, with more stable low dimensional representations has been observed in the prefrontal cortex of nonhuman primates (Druckmann and Chklovskii, 2012; Murray et al., 2017; Spaak et al., 2017; Sapountzis et al., 2022), and mice (such as the line attractor dynamics in Voitov and Mrsic-Flogel, 2022). The combination has been proposed as a mechanism for how the brain might encode working memory during the delay, while allowing for a stable readout by downstream regions. It is consistent with the notion of the PFC simultaneously allowing for the mixed selectivity that supports higher cognition (Fusi et al., 2016), while also playing a central role in reliably representing important memoranda in working memory for subsequent orchestration of action selection (Ridderinkhof et al., 2004; Howland et al., 2022; Soltani and Koechlin, 2022).

Of note, we found that time and the identity of the sample arm are both variables that are simultaneously encoded in the population activity in PFC. Importantly, they are ‘disentangled’, as they are represented in approximately orthogonal subspaces, so that a downstream linear readout can either read one variable or the other. Moreover, the time readout can easily generalize across remembered sample arms, and vice-versa, the sample arm readout can generalize across time. These kinds of representations are also called ‘abstract’, and are observed in NHP PFC (Bernardi et al., 2020; Fascianelli et al., 2024), and in other brain areas and species (Higgins et al., 2021; Nogueira et al., 2023; Boyle et al., 2024; Courellis et al., 2024).

### Mnemonic spatial representations in PFC delay activity reflect changes in entrained behavioral strategy

Comparative decoding analysis of retrospective and prospective spatial information in PFC delay activity revealed that mnemonic representations were most evident for the sample arm during trials in which the animal would subsequently choose to avoid returning to that sample arm (Figure 4). This suggests that in a large proportion of “error” trials, the animals would nonmatch to the sample arm not because they could not recall where they had been during the sample phase, but because either they could not remember the overall DMS task rule, or they were intentionally ignoring the known rule to employ a different behavioral strategy to explore the environment. Given their overall well-beyond-chance task performance, the consistency of the DMS rule across all training history, and the comparative absence of mnemonic representations on trials in which they followed a DMS rule, it is unlikely that the animals simply forgot the DMS task rule and thus failed to utilize evident retrospective representations for DMS adherence. Much more likely, then, is that the animals were intentionally shifting their behavioral rule in order to reassess environmental contingencies. In this light, many “error” trials are not conventional error trials that result from inattention or forgetting, but rather are reflections of an active behavioral strategy shift from exploitation of known environmental contingencies to exploration of less certain environmental contingencies. In this task, then, mnemonic representations were primarily present when animals deviated from the default behavioral strategy.

Might this mean, therefore, that mnemonic spatial WM representations in the PFC are some sort of foraging signal, present in general when the animal executes less repetitive behaviors (i.e. DNMS, but not DMS)? Such an interpretation is possible, but much less likely given the data from T-, Y-, and figure eight mazes (Jung et al., 1998; Ito et al., 2015; Spellman et al., 2015; Bolkan et al., 2017; Böhm and Lee, 2020; Vogel et al., 2022). Many of these tasks have a DNMS task design, and yet retrospective mnemonic spatial representations are rarely observed in correct trials, suggesting that mnemonic spatial WM representations in the PFC are not simply a reflection of general foraging or nonrepetitive behavior. If, instead, PFC mnemonic representations emerge to support a shift from the dominant behavioral strategy, one would expect such representations to be evident on error trials in these two-choice DNMS tasks as well—or at least the subset of them that result from intentional behavioral deviation, rather than inattention, interference, or forgetting. Perhaps because of the high baseline performance and high spatial interference on these two-choice tasks (Deacon and Rawlins, 2006; Missaire et al., 2017), a critical mass of intentional “error” trials is not commonly accrued or analyzed. Nevertheless, the general absence of mnemonic representations in correct trials in T-maze DNMS tasks and this RAM DMS task, and the presence of mnemonic representations in “error” trials in this RAM DMS task are consistent with a model of mnemonic PFC spatial WM representations that emerge to enable (or reflect) a deviation from the dominant behavioral strategy.

Putting the model together, within the framework of Baddeley and Hitch <u>(1974)</u>, working memory is instantiated via cooperative executive and mnemonic components. Given the cruciality of PFC activity to freely moving spatial working memory tasks (Vogel et al., 2022), the common absence of PFC mnemonic representations in such tasks suggests that the PFC may here be playing a primarily executive role, relying on non-prefrontal regions (Christophel et al., 2017; Sreenivasan and D’Esposito, 2019) for mnemonic representations. This representational characteristic seems to change, however, when the executive function demands increase, as in when the animal is seeking to deviate from its dominant behavioral strategy: in this context, mnemonic representations emerge locally within the PFC. Thus, within freely moving spatial working memory, the executive role of the PFC predominates, with mnemonic representations appearing locally in PFC mainly to support increased executive function load, thereby enabling greater neurobehavioral flexibility and a strategy change. This multicomponent essentiality of PFC activity for strategy-change neurobehavioral flexibility in spatial working memory is consistent with the essentiality of PFC activity for strategy-change neurobehavioral flexibility in other cognitive tasks, like the Wisconsin Card Sorting Task (Grant and Berg, 1948; Milner, 1963; Milner et al., 1964; Owen et al., 1991), and its analogues for nonhuman primates (Dias et al., 1996a, 1996b) and for rodents (Birrell and Brown, 2000; Bissonette et al., 2008; Benoit et al., 2022).

### Moving through the spatial environment may affect the nature of mnemonic representations

Spatial working memory tasks in which animals freely move and explore are appealing for understanding brain function because brains evolved to generate goal-directed movement (Krakauer et al., 2017; Cisek, 2022). As most of the spatial working memory literature involving freely-moving tasks is in rodents, the contributions of species differences versus tasks differences (versus other differences) in WM neural correlates cannot yet be resolved. WM neural correlates in PFC activity might differ in nonhuman primate visuospatial WM tasks and in rodent freely-moving spatial WM tasks not because of species differences, but because of freedom of movement differences. To further characterize the comparative contributions of task elements versus organismal elements to neural correlates, more freely-moving spatial WM experiments are needed in non-rodent species. Thankfully, a recent explosion of tool development for both freely-moving neural interrogation and behavioral analysis is making complex, freely-moving tasks ever more feasible across species (Kondo et al., 2018; Mathis et al., 2018, 2020; Juavinett et al., 2019; Luo et al., 2020; Mao et al., 2021; Van Daal et al., 2021; Liberti et al., 2022; Pereira et al., 2022; Zong et al., 2022; Forli and Yartsev, 2023; Guo et al., 2023; Klioutchnikov et al., 2023; Weinreb et al., 2023; Zhao et al., 2023).

